# “De novo Classification of Mouse B Cell Types using Surfaceome Proteotype Maps”

**DOI:** 10.1101/620344

**Authors:** Marc van Oostrum, Maik Müller, Fabian Klein, Roland Bruderer, Hui Zhang, Patrick G. A. Pedrioli, Lukas Reiter, Panagiotis Tsapogas, Antonius Rolink, Bernd Wollscheid

**Affiliations:** Biomedical Proteomics Platform, Department of Health Sciences and Technology, ETH Zurich, 8093 Zurich, Switzerland; Institute of Molecular Systems Biology at the Department of Biology, ETH Zurich, Zurich 8093, Switzerland; Biognosys AG, Schlieren, Switzerland; Developmental and Molecular Immunology; Department of Biomedicine; University of Basel; Basel; Switzerland; Department of Pathology, Johns Hopkins University, Baltimore, Maryland, USA

## Abstract

System-wide quantification of the cell surface proteotype and identification of extracellular glycosylation sites is challenging when sample is limiting. We miniaturized and automated the previously described Cell Surface Capture technology increasing sensitivity, reproducibility, and throughput. We used this technology, which we call autoCSC, to create population-specific surfaceome maps of developing mouse B cells and used targeted flow cytometry to uncover developmental cell subpopulations.

## Introduction

Cell types are typically identified and classified using selected cell-surface proteins as this sub-proteome reflects the maturity and functional state of a cell^1,2^. A comprehensive knowledge of the surface-exposed proteome (surfaceome) is necessary for classification of cell types and to enable linkage of distinct proteotypes to functional phenotypes. Global analysis of the surfaceome is challenging for technical reasons. High-throughput protein and RNA analyses are agnostic toward spatial information, and antibody-based technologies like flow or mass cytometry are limited in their multiplexing capabilities and by the availability of high-quality antibodies. Chemical biotinylation of surface glycoproteins has been used for systematic surfaceome interrogation by affinity purification of tagged proteins^3,4^ or peptides^5–7^ prior to mass spectrometry (MS) analysis, and protein-level enrichment methods that quantify peptides adjacent to a tagged glycopeptide have provided deep coverage of the plasma membrane proteome^3^. These methods, however, preclude *a priori* separation of enriched surface proteins from nonspecific background; instead, prior knowledge regarding surface localization is used to filter for known plasma membrane proteins. In contrast, Cell Surface Capture (CSC) technology enables direct identification of extracellular *N*-glycopeptides. Surface biotinylated *N*-glycopeptides are captured and enzymatically released by peptide:N-glycosidase F (PNGase F), which catalyses cleavage of asparagine-linked glycans. This leaves a deamidation within the NXS/T consensus sequence of formerly *N*-glycosylated peptides, indicating both surface localization and glycosylation site. The specific site information comes at cost of sensitivity, however. Hence CSC experiments are performed with up to 1 × 10^8^ cells per sample^6^. The large amounts of sample required mean that it is not practical to perform CSC on primary cells.

## Results

Miniaturization and automation have proven beneficial in phosphopeptide^8^, hydrazide^9^ and antibody-based^10^ enrichment processes. We reasoned that the same principles could be applied to the biotin-streptavidin system used within the CSC workflow. Additionally, we optimized the standard CSC technology, most prominently by inclusion of a catalyst for optimal surface labeling^11,12^. A schematic of the autoCSC workflow is shown in **Fig. 1a**. Briefly, live cells are oxidized under mild conditions with sodium-meta-periodate to generate aldehydes on cell-surface carbohydrates and are subsequently labelled with cell-impermeable biocytin-hydrazide in presence of catalyst 5-methoxyanthranilic acid^11,12^. After cell lysis and tryptic digestion, the resulting peptides are subjected to automated processing on a tailored liquid handling robot. Repeated aspiration through filter tips containing streptavidin resin provides a confined reaction space for efficient binding of biotinylated *N-*glycopeptides. After extensive washing, *N*-glycopeptides are released by PNGase F provided in a plate heated to 37 °C. Using high-resolution MS, labelled extracellular peptides are identified by deamidated asparagines within the NXS/T glycosylation consensus sequence resulting from PNGase F cleavage. SWATH-based data-independent acquisition (DIA) and targeted feature extraction are used to quantify surface-protein abundances across multiple conditions^13,14^.

First, we evaluated performance of the automated and miniaturized processing compared to the manual workflow. We prepared CSC-labelled peptides from approximately 6 × 10^8^ HeLa cells and distributed them over 30 samples containing about 5 mg peptide each. Twenty of these samples were processed in manual mode by two different researchers (Experimenters 1 and 2) and ten were subjected to automated processing. Experimenters 1 and 2 identified 325 and 290 extracellular *N*-glycopeptides, respectively (median), whereas the robot identified 1811 *N-*glycopeptides, a more than 5-fold increase over the manual process **(Fig. 1b)**. To assess quantitative reproducibility, we performed MS1-based label-free quantification and calculated coefficients of variation (CVs) for precursors that match *N*-glycopeptides. Automated processing showed the lowest variation with a median CV of 28 compared to 51 and 34 for Experimenter 1 and 2, respectively **(Fig. 1c)**. Thus, miniaturization and automation of the biotin-streptavidin enrichment resulted in increased sensitivity and higher quantitative reproducibility for the CSC workflow.

To evaluate the capability for multiplexed quantification of surfaceomes, we performed autoCSC with 11 commonly used cancer cell lines. After cell labelling and digestion, samples containing approximately 1 mg peptide were processed, and *N-*glycoproteins were quantified with DIA-MS. We quantified 1,697 unique glycosylated asparagines located within proteotypic peptides from 900 protein groups with a median of two sites per protein group **(Supplementary Fig. 1, Supplementary Table 1)**. Of these, 192 glycosylation sites were previously unknown based on Uniprot database annotation **(Supplementary Fig. 2a)**. The median per cell type was 602 surface proteins, two-fold higher compared to 301 reported in the largest CSC data repository, the Cell Surface Protein Atlas^6^, despite using 5-to 33-fold lower amounts of input peptides. Hierarchical clustering and principal component analysis grouped samples originating from the same cell type in close proximity **(Fig. 1d** and **Fig. 1e)**, demonstrating that autoCSC can reliably subclassify cell types based on surfaceome.

We hypothesized that autoCSC technology would enable phenotyping of freshly isolated primary cell populations. B cell development consists of successive cellular stages beginning in the bone marrow and continuing in peripheral lymphoid tissues^14,15^. Developing B cells are particularly challenging to characterize by CSC due to limited sample availability from mice, relatively small size, and little diversity in surfaceome composition^15^. In order to probe applicability of autoCSC, we prepared a dilution series from 30 to 0.1 × 10^6^ cells of a B lymphoma cell line and quantified *N*-glycoproteins. Compared to the number of surface proteins identified from a sample of 30 × 10^6^ cells, we recovered a median of 62% with 1 × 10^6^, although the variance increased below 5 × 10^6^ **(Supplementary Fig. 3)**. With this limitation in mind, we set out to *de novo* map and quantitatively compare the surfaceomes of developing B cells using autoCSC. From mice, we isolated nine consecutive stages of B cell populations **(Fig. 2a)** with a maximum of 1 × 10^6^ cells per sample using fluorescence-activated cell sorting (FACS). With autoCSC, we quantified 248 unique glycosylated asparagines located within proteotypic peptides and grouped them into 147 protein groups with a median of two unique sites per protein group **(Fig. 2b and Supplementary Table 2)**. Of these, 25 glycosylation sites were previously unknown based on Uniprot database annotation **(Supplementary Fig. 2b)**. Some proteins were found exclusively on a particular population, for example CD80 and CD130 on B1 cells **(Supplementary Figure 5)** and are potentially useful as stage-specific markers **(Fig. 2b)**. Furthermore, we identified several surface proteins that were either absent or of considerably lower abundance in the bone marrow until the immature B stage, but showed stable surface abundance in later stages **(Fig. 2b-f and Supplementary Fig. 5)**. We hypothesized that these could be used to further split the immature B stage into subpopulations with different maturities. Therefore, we performed flow cytometry analysis to identify differences in surface abundance among individual cells within each population **(Supplementary Fig. 6)**. First, we asked whether flow cytometry reproduced the autoCSC results for the selected proteins. All showed a strong positive correlation with an average Spearman’s rho of 0.77, indicating very good agreement between the two methods **(Fig. 2d)**. As a reference we retrieved RNA-seq data for the corresponding populations from the ImmGen database and calculated correlation coefficients with flow cytometry **(Fig. 2d)**^16^**. Interestingly, flow cytometry revealed a bimodal distribution of CD20 within the immature B subpopulation (Supplementary Fig. 6)**. For CD180, we found a broad distribution covering more than two orders of magnitude **(Supplementary Fig. 6)**. Co-staining for CD20 and CD180 across all populations revealed that prior to CD180 upregulation, CD20 abundance increased during development **(Fig. 2g)**. Based on our data, we were able to further divide the immature B population into three subpopulations: double negative, positive for CD20 but not CD180, and positive for both CD20 and CD180 **(Fig. 2h)**. We then asked whether the three subpopulations are different regarding maturity and found that the double-positive population had a significantly higher number of IgD^+^ cells **(Fig. 2i)** accompanied by higher surface abundance of BAFF receptor **(Fig. 2j)** compared to single-positive and double-negative subpopulations, indicating that the double-positive subpopulation is at the verge of initiating migration from bone marrow to the spleen.

## Discussion

In summary, miniaturized and automated CSC technology was validated and used to analyse cancer cell lines and developing primary B cells. mRNA abundance is not sufficient to infer protein levels in many scenarios^17^, and to predict genotype-phenotype relationships it is necessary to go beyond global protein levels and determine proteoform abundances within the spatial domains where activity is required for function^18^. CSC technology utilizes an amino acid modification within a consensus sequence that reports on an initial cell-surface-restricted labelling event, enabling specific quantification with subcellular resolution. Direct identification of tagged sites has proven beneficial for spatial proteomic workflows^19^ but requires the detection of a peptide bearing the modified amino acid. This caveat makes it particularly challenging to achieve identification of large numbers of formerly glycosylated sites when sample amounts are low. Insufficient sensitivity resulting at least partially from manual processing has limited CSC to specimens that can be produced in large quantities in the range of 50 - 100 Mio cells per sample. Physical confinement of the reaction space and automation of the biotin-streptavidin system within the autoCSC technology bridged this gap and reached the sensitivity and throughput required for broad application of surfaceome screening with primary cellular material. These improvements will enable surfaceome proteotype maps of primary cell samples for translational research applications and systems-scale surfaceome research.

**Figure 1.**
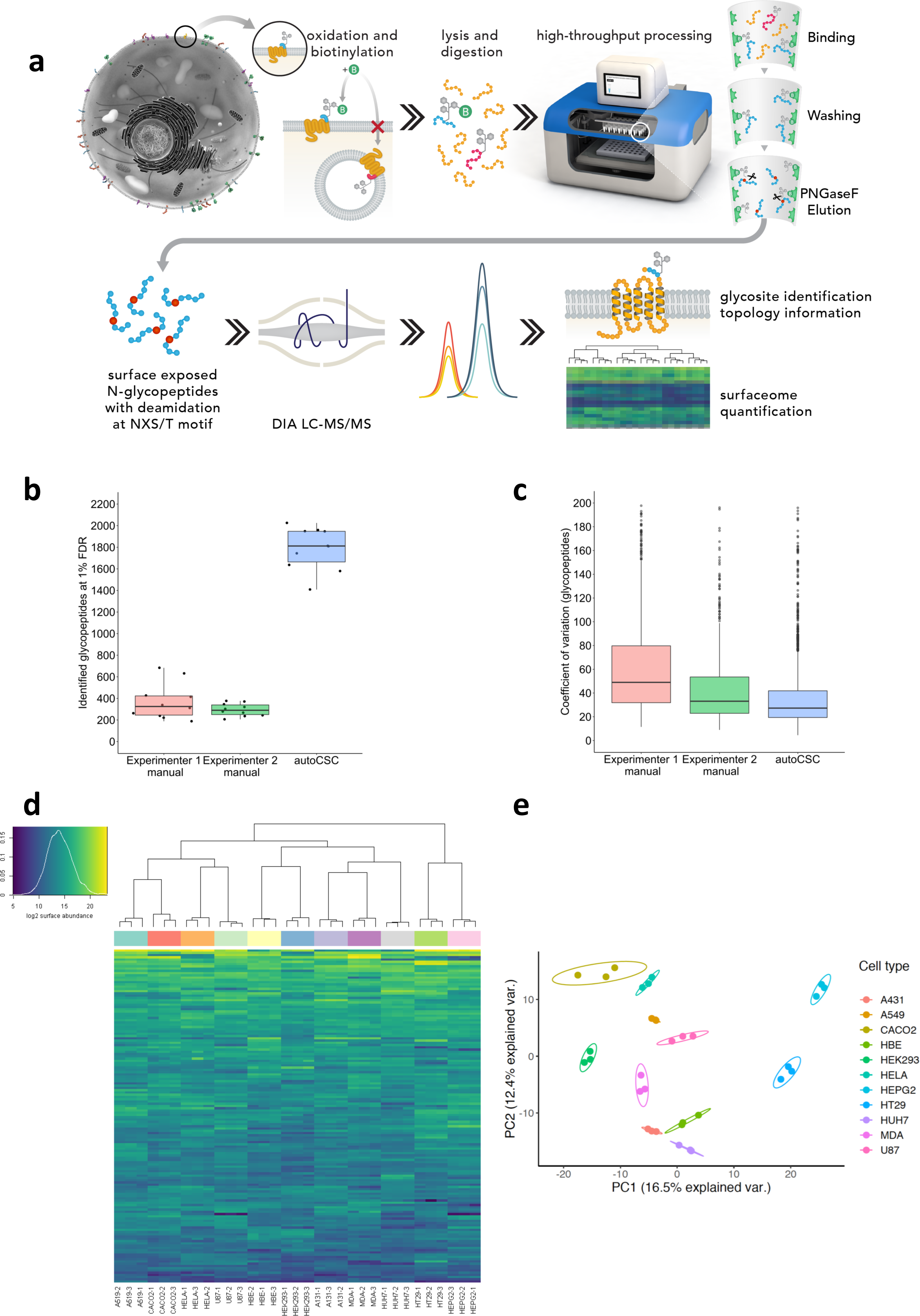

**Figure 2.**
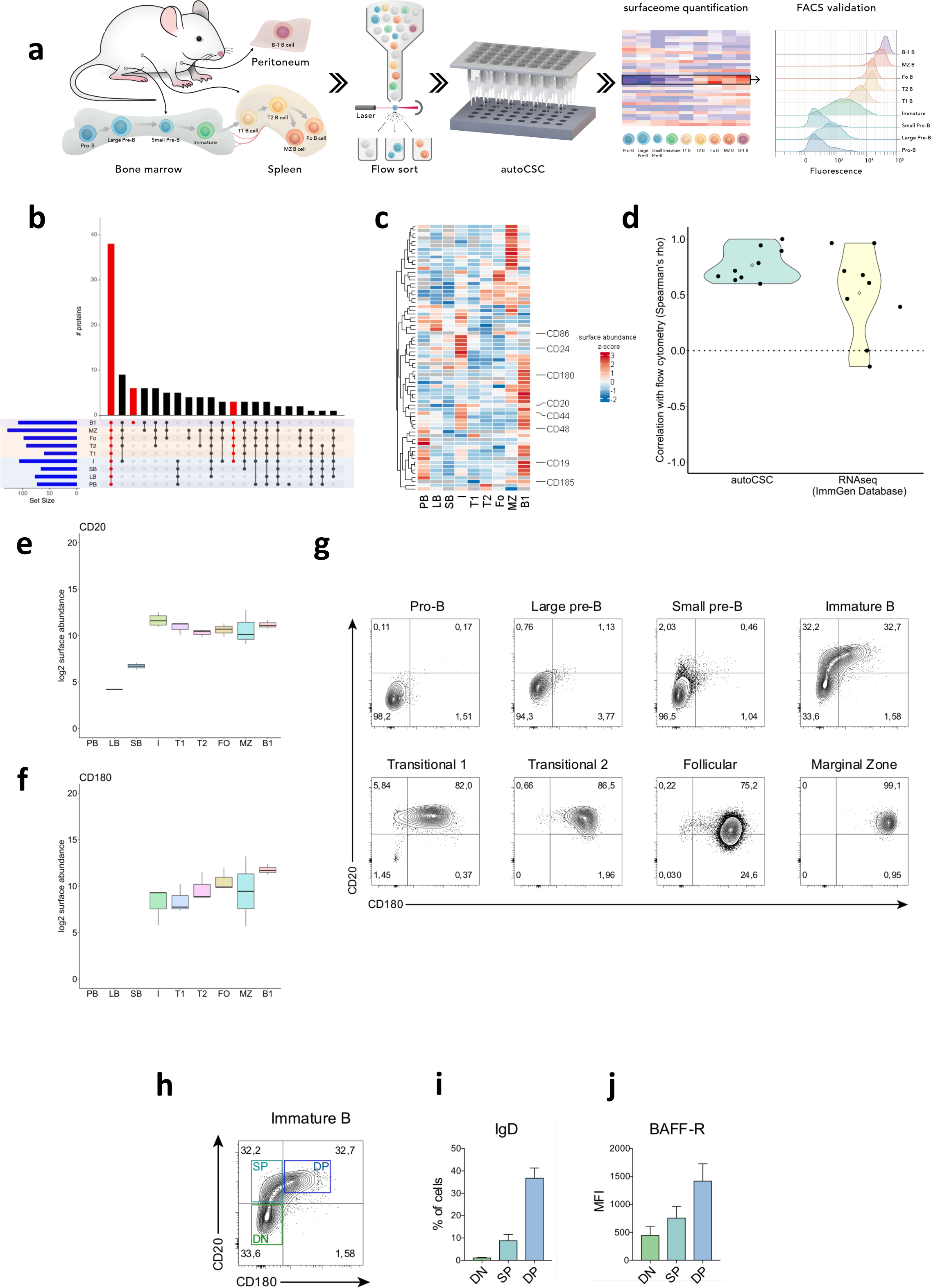

## Methods

### Chemicals

All chemicals were purchased from Sigma unless stated otherwise.

### Mice

C57BL/6 mice were bred and maintained in our animal facility in pathogen-free conditions. All mice used were 5-7 weeks old. All animal experiments were carried out under institutional guidelines (authorization number 1888 from canton Basel-Stadt veterinary office).

### Cell culture

All cell lines were purchased from ATCC, except 16HBE14o-, which was a kind gift from Jason Mercer (MRC-Laboratory for Molecular Cell Biology, University College London). Cells were grown at 37 °C and 5% ambient CO_2_. HepG2, HEK-293, HeLa Kyoto, Caco-2, U87, A431, MDA-MB-231, Huh-7, and A549 cells were cultured in Dulbecco’s Modified Eagle’s Medium, high glucose, GlutaMAX supplement (DMEM, Thermo Scientific), 10% fetal bovine serum (FBS), and 1% penicillin-streptomycin (PS). 16HBE14o- and SU-DHL-6 cells were grown in RPMI 1640 with 1.5 mM GlutaMAX, 1% PS, and 10% FBS. HT29 cells were cultured in McCoy′s 5a medium with 1.5 mM GlutaMAX, 10% FBS, and 1% PS.

### Antibodies, flow cytometry, and sorting

Cells were flushed from femurs and tibias of the two hind legs and from the peritoneum of the mice or single-cell suspensions of spleen cells were made. Staining was performed in PBS containing 0.5% BSA and 5 mM EDTA. The following antibodies were used for flow cytometry (from BD Biosciences, eBioscience, BioLegend, or produced in house): anti-CD117 (2B8), anti-CD19 (1D3), anti-CD127 (SB/199), anti-IgM (M41), anti-IgD (1.19), anti-CD93 (PB493), anti-CD11b (M1.7015), anti-CD5 (53-7.3) anti-CD23 (B3B4), anti-CD44 (IM7), anti-CD48 (HM48-1), anti-CD24 (M1/69), anti-CD20 (SA275A11), anti-CD180 (RP/14), anti-CD150 (TC15-12F12.2), anti-CXCR5 (2G8), anti-PD-L1 (10F.9G2)), anti-CD130 (4H1B35), anti-CD80 (16-10A1), anti-CD86 (GL1), an anti-mBAFF-R (9B9). For flow cytometry a BD LSRFortessa (BD Biosciences) was used, and data were analysed using FlowJo Software (Treestar). For cell sorting, a FACSAria IIu (BD Biosciences) was used (>98% purity). For autoCSC, sorting was performed on four different days generating four biological replicates. Two technical replicates with 1 × 10^6^ cells per biological replicate were processed for B cell populations (SB, I, FO, MZ) with sorting yields of 2 × 10^6^ cells.

### Cell Surface Capture - labeling and digestion

#### Labelling and digestion for CSC

Surface glycoproteins on live cells were gently oxidized with 2 mM NaIO_4_ (20 min, 4 °C) in labelling Buffer (LB) consisting of phosphate-buffered saline, pH 6.5. Cells were washed once in LB and subsequently biotinylated in LB containing 5 mM biocytin hydrazide (Pitsch Nucleic Acids, Switzerland) and 5 mM 5-methoxyanthranilic acid for 1 h at 4 °C min. Cells were washed three times with LB and harvested, lysed in lysis buffer (100 mM Tris, 1% sodium deoxycholate, 10 mM TCEP, 15 mM 2-chloroacetamide) by repeated sonication using a VialTweeter (Hielscher Ultrasonics), and heated to 95 °C for 5 min. Proteins were digested with trypsin overnight at 37 °C using an enzyme-to-protein ratio of 1:50. For the CSC dilution series and experiments with B cells, LB used for washes contained 5% FBS, samples were pre-incubated with buffer containing FBS prior to lysis, and digestion was done using LysC (Wako) and sequencing-grade trypsin (Promega), both with an enzyme-to-protein ratio of 1:200. In order to inactivate trypsin and precipitate deoxycholate, samples were boiled for 20 min, acidified with 10% formic acid to approximately pH 3, and centrifuged 10 min at 16,000 g. Peptide concentrations were determined in the supernatant using a NanoDrop 2000 instrument (Thermo Scientific). For experiments where the cell numbers were not normalized prior CSC labeling, the peptide mixtures were normalized before aliquoting into a 96-well sample plate for automated processing.

### Automated processing

A Versette liquid handling robot (Thermo Scientific) was equipped with a Peltier element in order to adjust temperature within 96-well plates during glycopeptide elution. Streptavidin tips were prepared by pushing a bottom filter membrane into disposable automation tips (Thermo Scientific). Each tip was filled with 80 µl of Pierce Streptavidin Plus UltraLink Resin (Thermo Scientific), and tips were sealed by compressing the resin with a top filter membrane. Assembled tips were attached to the liquid handling robot and washed with 50 mM ammonium bicarbonate by repeated cycles of aspiration and dispensing (mixing). Likewise, biotinylated peptides were bound to the streptavidin resin over 2.5 hours of mixing cycles. Subsequently the streptavidin tips were sequentially washed with 5 M NaCl, StimLys Buffer (100 mM NaCl, 100 mM glycerol, 50 mM Tris, 1% Triton X-100), with 50 mM ammonium bicarbonate, 100 mM NaHCO_3_, pH 11, and with 50 mM ammonium bicarbonate. For glycopeptide elution, streptavidin tips were incubated overnight in 50 mM ammonium bicarbonate containing 1,000 units PNGase F (New England Biolabs) at 37 °C. The sample plate was then removed from the liquid handling robot and acidified to pH 2–3 with formic acid. Peptides were desalted with C18 UltraMicroSpin columns (The Nest Group) according to the manufacturer’s instructions and dried in a SpeedVac concentrator (Thermo Scientific). For comparison of the automated with the manual workflow, manual processing was done analogously: Biotinylated peptides were bound to streptavidin in 1.5-mL tubes by addition of 80 μL washed Streptavidin Plus UltraLink Resin (Pierce), and samples were incubated for 2.5 hours on a slow rotator. Beads were washed in Mobicols (Boca Scientific) connected to a Vac-Man Laboratory Vacuum Manifold (Promega) using the same buffers. Washed beads were incubated with 50 mM ammonium bicarbonate containing 1,000 units PNGase F (New England Biolabs) in a head-over-head rotator overnight at 37 °C.

### Liquid chromatography–tandem mass spectrometry (LC-MS/MS) analysis

For MS analysis, peptides were reconstituted in 5% acetonitrile and 0.1% formic acid containing iRT peptides (Biognosys). Depending on instrument availability and estimated required sensitivity, experiments were analysed on different LC-MS/MS systems.

The peptides resulting from the comparison between automated and manual processing were analysed in DDA mode. They were separated by reverse-phase chromatography on a high-pressure liquid chromatography (HPLC) column (75-μm inner diameter; New Objective) packed in-house with a 15-cm stationary phase ReproSil-Pur 120A C18-AQ 1.9 µm (Dr. Maisch GmbH) and connected to an EASY-nLC 1000 instrument equipped with an autosampler (Thermo Scientific). The HPLC was coupled to a Q Exactive plus mass spectrometer equipped with a nanoelectrospray ion source (Thermo Scientific). Peptides were loaded onto the column with 100% buffer A (99% H_2_O, 0.1% formic acid) and eluted at a constant flow rate of 300 nL/min for 50 min with a segmented gradient from 6 to 35% buffer B (99.9% acetonitrile, 0.1% formic acid). Mass spectra were acquired in a data-dependent manner (top 12). In a standard method for medium-to-low abundance samples, high-resolution MS1 spectra were acquired at 70,000 resolution (automatic gain control target value 3 × 10^6^) to monitor peptide ions in the mass range of 375–1500 m/z, followed by high-energy collisional dissociation (HCD)-MS/MS scans at 35,000 resolution (automatic gain control target value 1 × 10^6^). To avoid multiple scans of dominant ions, the precursor ion masses of scanned ions were dynamically excluded from MS/MS analysis for 30 s. Single-charged ions and ions with unassigned charge states or charge states above 6 were excluded from MS/MS fragmentation.

The peptides resulting from the analysis of 11 common cell lines and the SU-DHL-6 dilution series were analysed in DIA and DDA modes for spectral library generation. For spectral library generation, a fraction of samples originating from the same cell line were pooled to generate 11 mixed pools. For the SU-DHL-6 dilution series, a pooled sample was generated for library generation by mixing fractions of the replicates with more than 10 × 10^6^ cells input. Each sample was separated using a self-packed analytical PicoFrit column (75 µm × 50 cm length) (New Objective) packed with ReproSil-Pur 120A C18-AQ 1.9 µm (Dr. Maisch GmbH) using an EASY-nLC 1200 (Thermo Scientific). Peptides were loaded onto the column with 100% buffer A (99% H_2_O, 0.1% formic acid) and eluted at a flow rate of 250 nL/min with a segmented gradient from 1 to 52% buffer B (85% acetonitrile, 0.1% formic acid). Data were acquired on a Q Exactive HF-X mass spectrometer (Thermo Scientific). The DIA method contained 20 DIA segments of 30,000 resolution with IT set to auto, AGC of 3×10^6^, and a survey scan of 120,000 resolution with 60 ms max IT and AGC of 3×10^6^. The mass range was set to 350-1650 m/z. The default charge state was set to 3. Loop count 1 and normalized collision energy stepped at 25.5, 27, and 30. For the DDA, a TOP10 method was recorded with 60,000 resolution of the MS1 scan and 20 ms max IT and AGC of 3×10^6^. The MS2 scan was recorded with 60,000 resolution of the MS1 scan and 110 ms max IT and AGC of 3×10^6^. The covered mass range was identical to the DIA. The isolation width was set to 8 Th and the normalized collision energy to 27%.

The peptides resulting from the B cell populations were analysed in DIA and DDA modes for spectral library generation. For spectral library generation, a fraction of the samples originating from the same B cell population were pooled to generate 13 mixed pools, additionally we included 18 CSC samples from unsorted single cell suspensions from spleen (13) and bone marrow (5). Each sample was separated using an Acclaim PepMap RSLC C18, 250 mm length, 75 µm inner diameter, 2 µm particle size (Thermo Scientific), connected to a stainless steel emitter (Thermo Scientific, part. no. ES 542) using an EASY-nLC 1200 (Thermo Scientific). Peptides were loaded onto the column with 100% buffer A (98% H_2_O, 2% acetonitrile, 0.15% formic acid) and eluted at a flow rate of 250 nL/min with a segmented gradient from 1 to 38% buffer B (80% acetonitrile, 0.15% formic acid). Data were acquired on a Fusion Lumos mass spectrometer (Thermo Scientific). The DIA method contained 18 DIA segments of 30,000 resolution with IT set to 50 ms and AGC of 5×10^5^ and a survey scan of 60,000 resolution with 180 ms max IT and AGC of 1×10^6^. The mass range was set to 350-1650 m/z. The default charge state was set to 2. Loop count 1 and normalized collision energy stepped by 2% at 27.5%. For the DDA, MS1 scans were recorded with 60,000 resolution and 50 ms max IT and AGC of 1×10^6^. With a 3 s cycle time, MS2 scans were recorded with 30,000 resolution and 120 ms max IT and AGC of 2×10^5^. The covered mass range was identical to the DIA. The isolation width was set to 1.4 Th, and the normalized collision energy was stepped by 2% at 27.5%. All mass spectrometric data and acquisition information were deposited to the ProteomeXchange Consortium (www.proteomexchange.org/) via the PRIDE partner repository (data set identifier: PXD013627).

### Data analysis DDA LC-MS/MS

The peptides resulting from the comparison between automated and manual processing were analysed in DDA mode. RAW data were converted to mzML using MSconvert. Fragment ion spectra were searched with COMET (v27.0) against UniprotKB (Swiss-Prot, Homo sapiens, retrieved April 2018) containing common contaminants and MS standards. The precursor mass tolerance was set to 20 ppm. Carbamidomethylation was set as a fixed modification for cysteine, oxidation of methionine and deamidation of arginine were set as variable modifications. Probability scoring was done with PeptideProphet of the Trans-Proteomic Pipeline (v4.6.2). Peptide identifications were filtered for a FDR of ≤1% and presence of consensus NXS/T sequence and deamidation (+0.98 Da) at asparagines. Non-conflicting peptide feature intensities were extracted with Progenesis QI (Nonlinear Dynamics) for label-free quantification and determination of CVs.

### Data analysis DIA LC-MS/MS

LC-MS/MS DIA runs were analysed with Spectronaut Pulsar × version 12 (Biognosys)^14,20^ using default settings. Briefly, a spectral library was generated from pooled samples measured in DDA (details above). The collected DDA spectra were searched against UniprotKB (Swiss-Prot, Homo sapiens or Mus musculus, retrieved April 2018) using the Sequest HT search engine within Thermo Proteome Discoverer version 2.1 (Thermo Scientific). We allowed up to two missed cleavages and semi-specific tryptic digestion. Carbamidomethylation was set as a fixed modification for cysteine, oxidation of methionine and deamidation of arginine were set as variable modifications. Monoisotopic peptide tolerance was set to 10 ppm, and fragment mass tolerance was set to 0.02 Da. The identified proteins were assessed using Percolator and filtered using the high peptide confidence setting in Protein Discoverer. Analysis results were then imported to Spectronaut Pulsar version 12 (Biognosys) for the generation of spectral libraries.

Targeted data extraction of DIA-MS acquisitions was performed with Spectronaut version 12 (Biognosys AG) with default settings using the generated spectral libraries as previously described^14,20^. The proteotypicity filter “only protein group specific” was applied. Extracted features were exported from Spectronaut using “Quantification Data Filtering” for statistical analysis with MSstats^21^ (version 3.8.6) using default settings. Briefly, features were filtered for use for calculation of Protein Group Quantity as defined in Spectronaut settings, common contaminants were excluded, and presence of consensus NXS/T sequence including a deamidation (+0.98 Da) at asparagine was required. Features were then log-transformed, normalized, and quantified as protein abundance for each sample and condition using the *quantification* function in MSstats^21^. Protein abundance per sample or conditions was used for further analysis and plotting. autoCSC experiments comparing 11 cell lines and developing B cells were performed in multiple replicates (quadruplicate biological replicates and where possible additional technical replicates). Technical replicates per biological replicate were consolidated in MSstats. Outliers were removed to retain minimally three biological replicates, generating 31 samples for B cell populations and 33 samples for the 11-cell lines comparison for final quantification.

### Correlation analysis

Proteins with abundance values (non-NA) for flow cytometry and CSC for at least four populations were considered for analysis (CD20, CD24, PDL1, CD180, CD44, CD48, CD150, CD19, CXCR5). If one condition only was not applicable it was set as zero assuming an abundance value below limit of detection. Spearman correlation coefficients were calculated and plotted using R. RNA-seq data were retrieved for the same proteins from the ImmGen database (www.immgen.org). In the retrieved dataset the following populations were sorted based on the same markers as our flow cytometry analysis: Transitional 1, Transitional 2, Follicular, and Marginal Zone. The following populations were considered equivalent between ImmGen and our flow cytometry analysis: Pro-B to FrB/C, Immature B to FrE, and B1 to B1b. Large and small pre-B are not included in ImmGen database and were therefore not included in the correlation analysis between flow cytometry and RNA.

### Glycosylation site analysis

For glycosylation site counting the following rules were followed: i) only glycosylated peptides conforming to the NX[STC] consensus sequence were considered; ii) to avoid inflating the count, non-proteotypic peptides were arbitrarily assigned to a single protein in the protein group; iii) if a glycosylated peptide could be mapped to multiple positions within the same protein, both positions were kept, unless one of the mappings resulted in a higher number of sites matching the consensus NX[STC] motif, in which case only this one was kept. When comparing the glycosylation sites identified in this study to the ones annotated in UniProt, only proteins identified in this study as having at least one glycosylation site were considered in UniProt.

## Author contributions

M.v.O. and M.M. performed all experiments except those noted below. F.K. and P.T. performed flow cytometry and FACS. M.v.O., M.M., R.B., L.R. and B.W. optimized and performed LC-MS/MS acquisition. M.v.O., M.M., F.K., P.T., R.B. and P.G.A.P. analyzed data. P.G.A.P. performed glycosylation site analysis. H.Z. contributed new analytical tools. M.v.O., M.M., F.K., P.T., A.R. and B.W. designed research. M.v.O., M.M. and B.W. conceived the project and wrote and revised the manuscript.

## Acknowledgements

We acknowledge A. Leitner & R. Aebersold for sharing and maintaining their instrumentation. We are grateful to the members of the R.A. and B.W. research groups for suggestions and support at all stages of the project. We thank K. Novy for facilitating collaboration, J. R. Wyatt for text editing, J. Niebel and S. Sung for technical and T. Splettstoeser for graphical support. This work benefited from data assembled by the Immgen consortium. The following agencies are thanked for funding: ETH (grant ETH-30 17-1 and grant ETH-25 15-2) and Swiss National Science Foundation (grant 31003A_160259) for B.W.. A.R. was holder of a chair in immunology endowed by L. Hoffman-La Roche Ltd, Basel.

## Supplementary Figure Legends

**Supplementary Figure 1.**
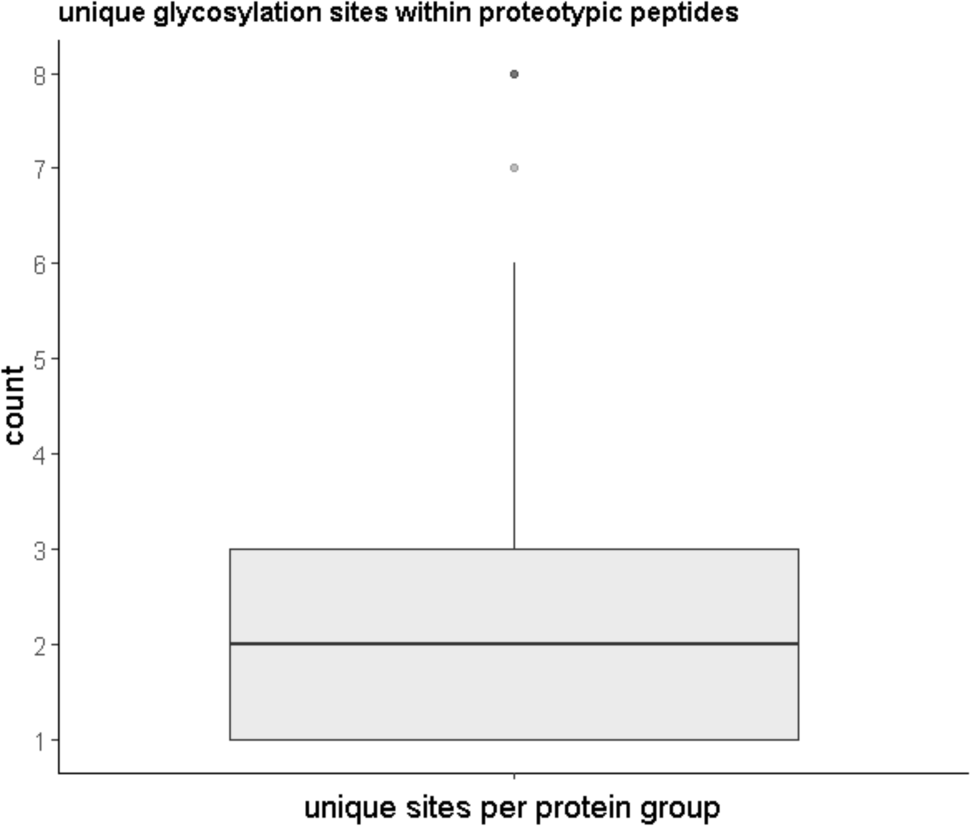
Number of identified glycosylation site per protein group. Distribution of unique glycosylation sites within proteotypic peptides per quantified protein group.

**Supplementary Figure 2.**
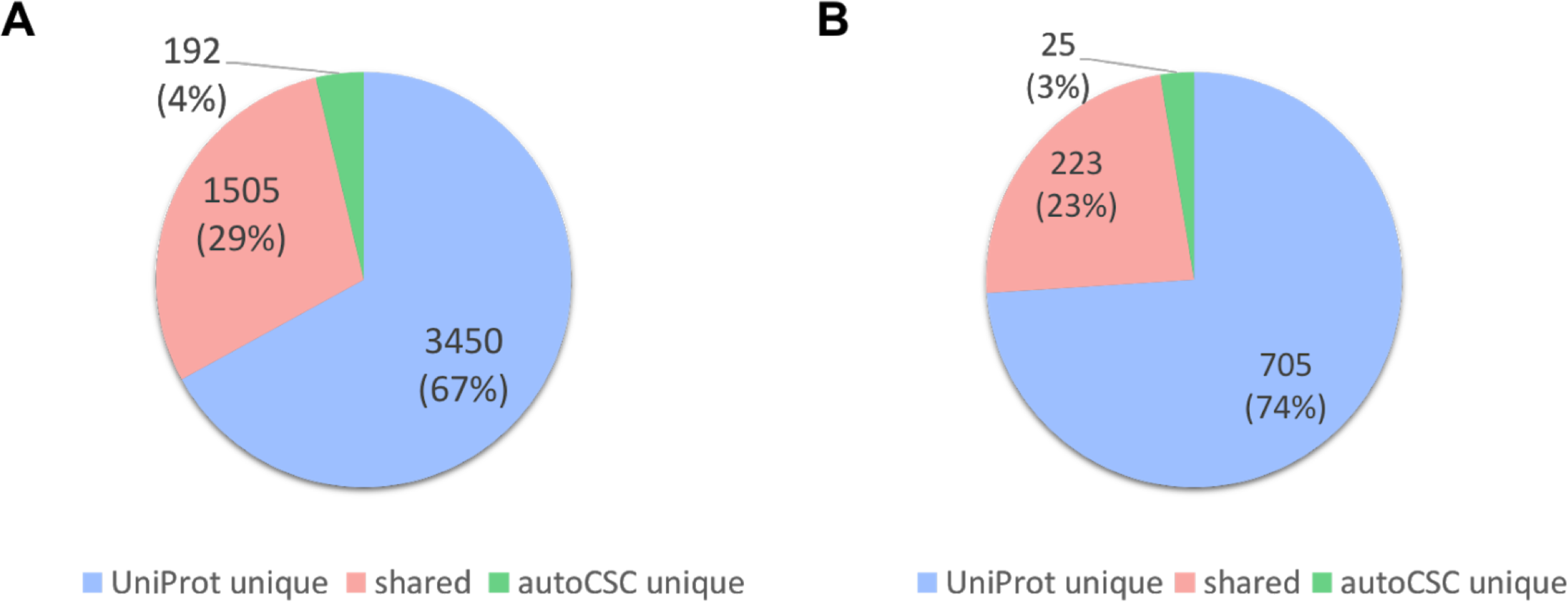
Number of glycosylation sites that are either annotated by Uniprot, identified by autoCSC or both in the surfaceome dataset of a) 11 common human cell lines or b) primary sorted mouse B-cells.

**Supplementary Figure 3.**
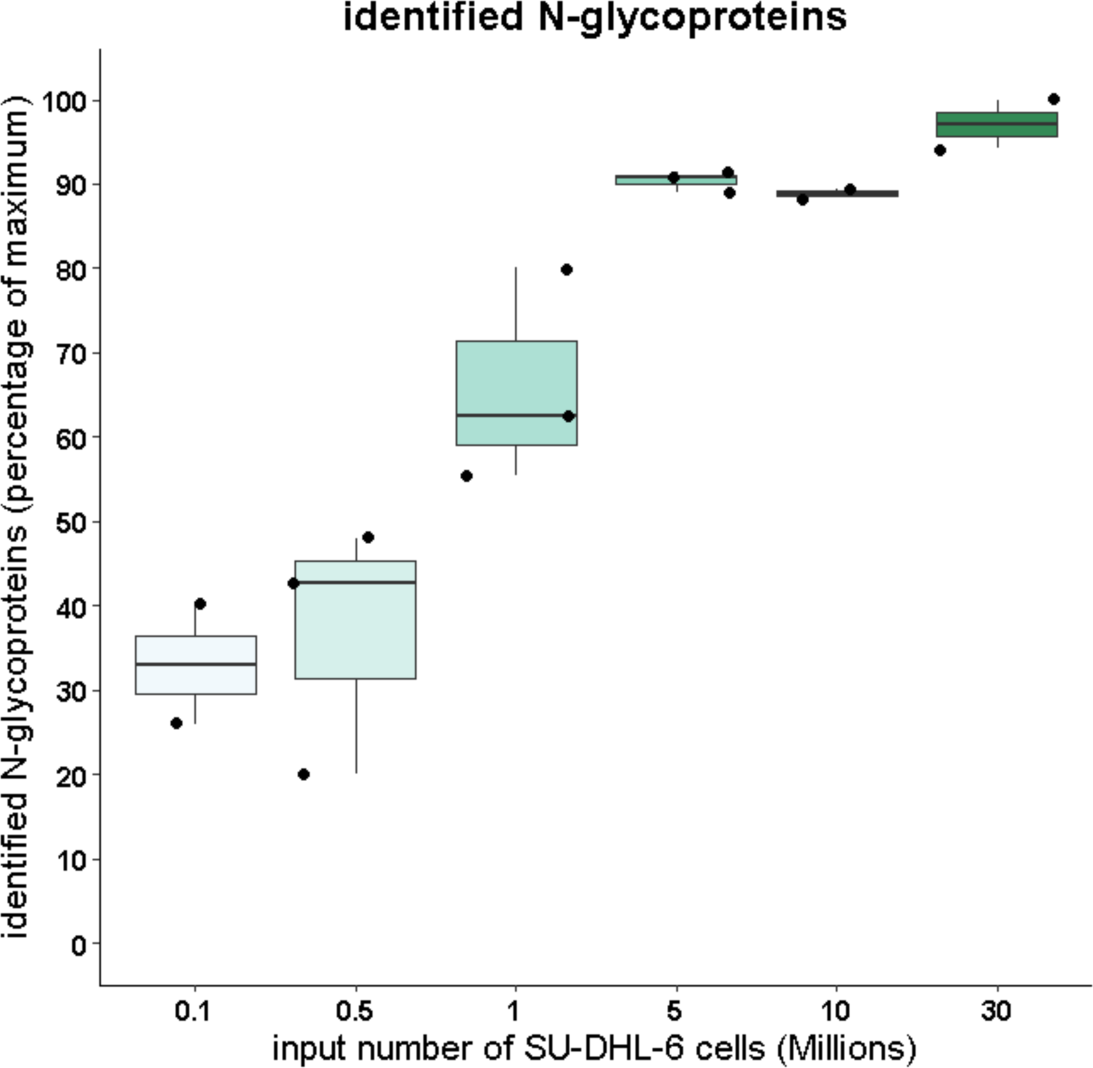
Dilution series of cell numbers used for autoCSC. Quantified *N*-glycopeptides using 0.1 to 30 × 10^6^ SU-DHL-6 cells as input for autoCSC.

**Supplementary Figure 4.**
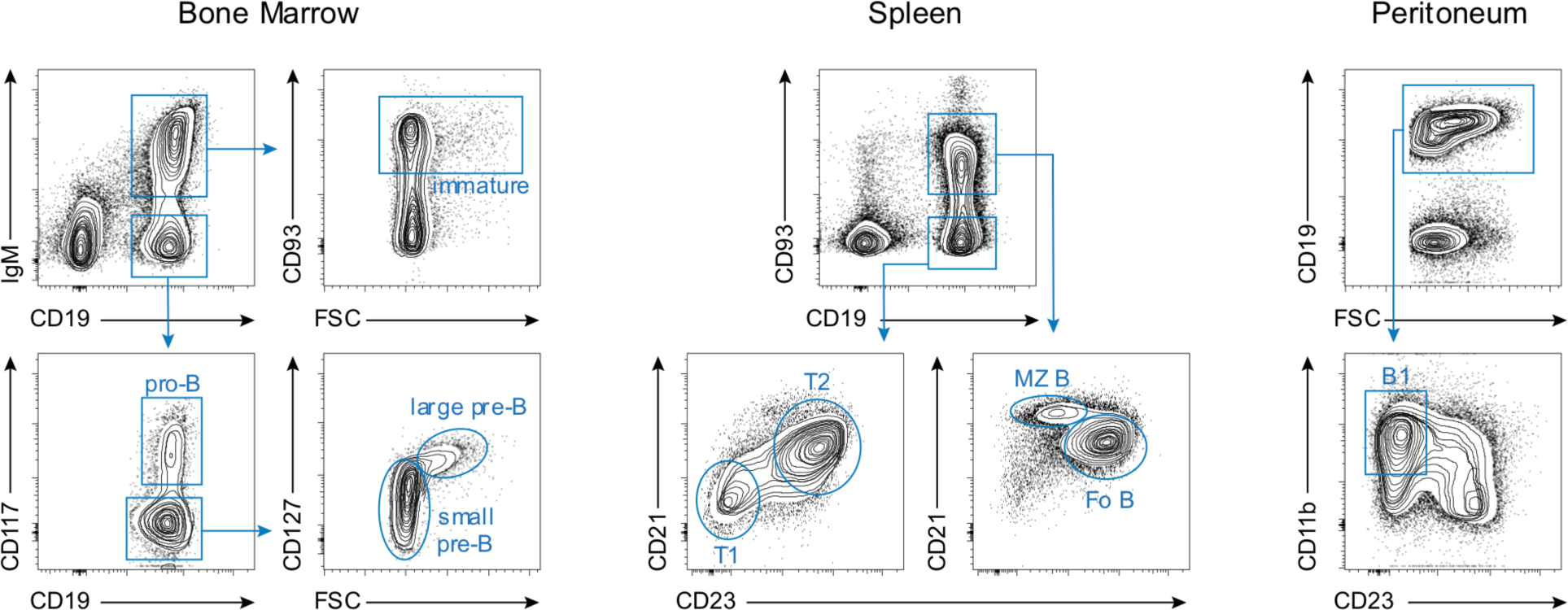
FACS strategy for mouse B cell populations. From the bone marrow: proB, CD19^+^IgM^-^CD117^+^; large-preB, CD19^+^IgM^-^CD117^-^CD127^+^FSC^large^; small-preB, CD19^+^IgM^-^CD117^-^CD127^-^FSC^small^; and immature B, CD19^+^IgM^+^CD93^+^. From the peritoneum: B1, CD19^+^CD23^-^CD11b^+^. From the spleen: Transitional-1, CD19^+^CD93^+^CD23^-^CD21^-^; Transitional-2, CD19^+^CD93^+^CD23^+^CD21^+^; Follicular (Fo), CD19^+^CD93^-^CD23^+^CD21^low^; and Marginal Zone (MZ), CD19^+^CD93^-^CD23^low^CD21^+^.

**Supplementary Figure 5.**
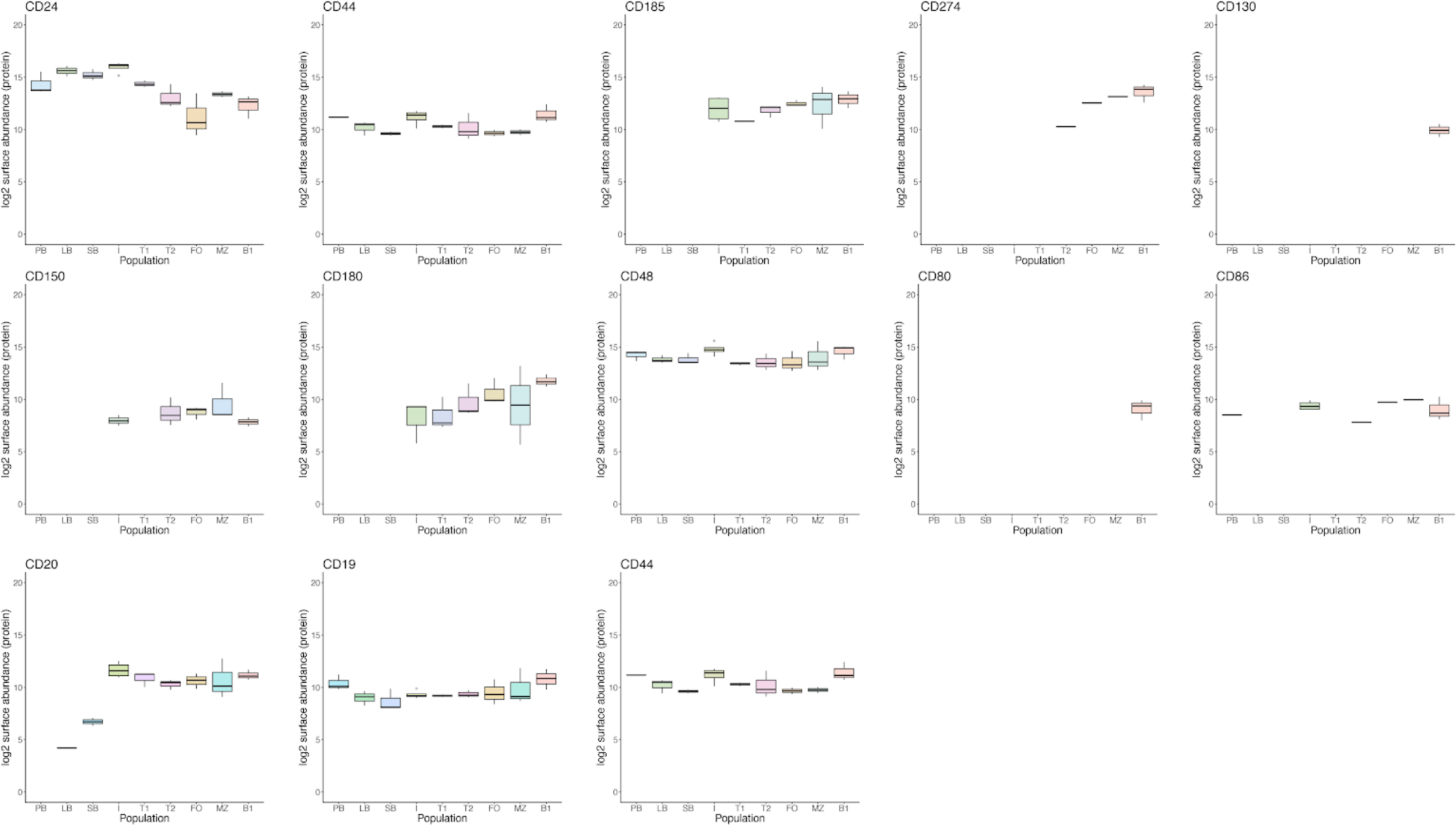
| Boxplots of surface abundances across B cell populations for proteins selected for follow-up flow cytometry analysis.

**Supplementary Figure 6.**
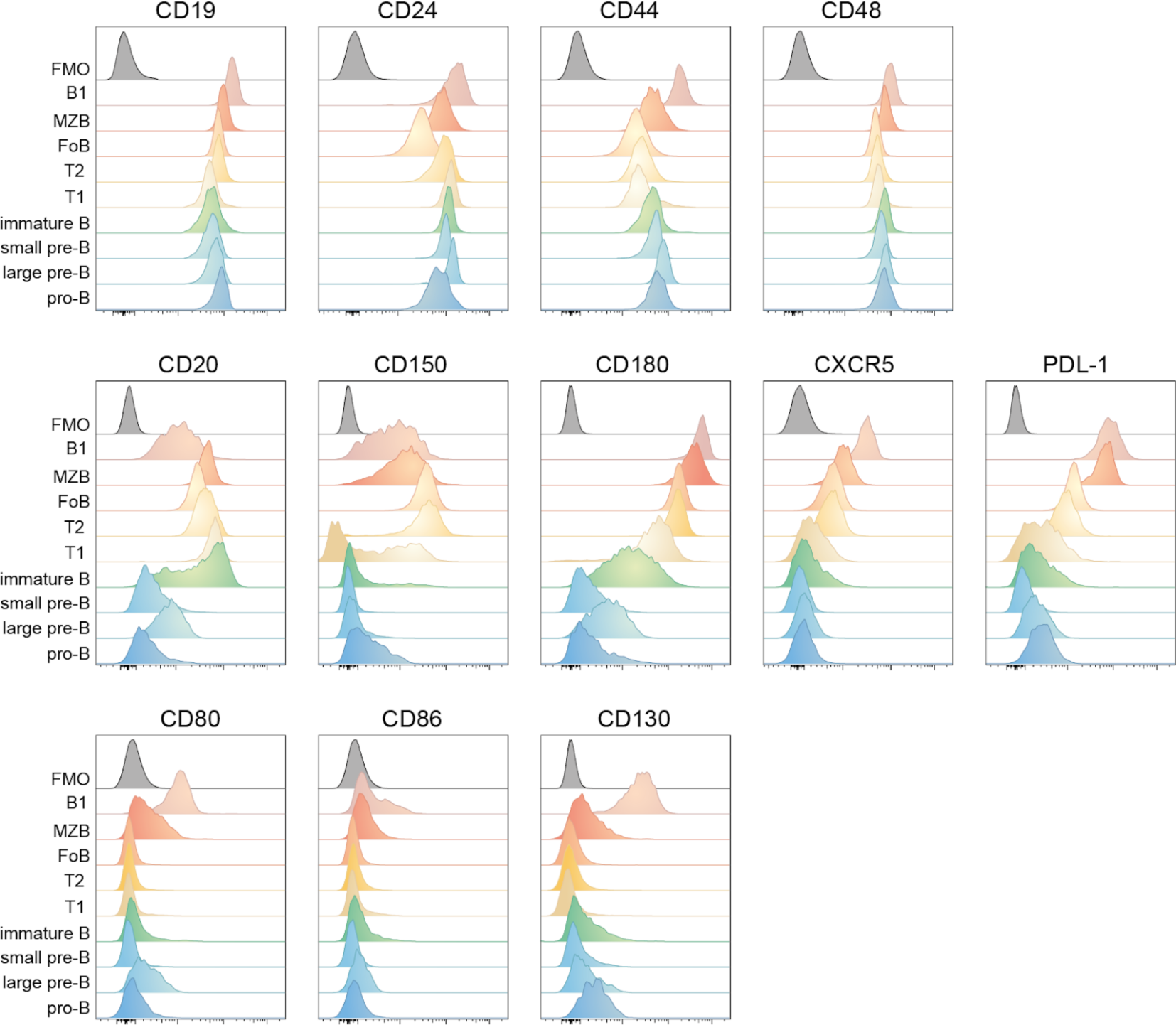
FACS plots of developing B cell populations for proteins selected from B cell surfaceomes generated by autoCSC.

## References

1. Orre, L. M. et al. SubCellBarCode: Proteome-wide Mapping of Protein Localization and Relocalization. Mol. Cell 73, 166–182.e7 (2019).

2. Itzhak, D. N. et al. A Mass Spectrometry-Based Approach for Mapping Protein Subcellular Localization Reveals the Spatial Proteome of Mouse Primary Neurons. Cell Rep. 20, 2706–2718 (2017).

3. Kalxdorf, M., Gade, S., Eberl, H. C. & Bantscheff, M. Monitoring Cell-surface N-Glycoproteome Dynamics by Quantitative Proteomics Reveals Mechanistic Insights into Macrophage Differentiation. Mol. Cell. Proteomics 16, 770–785 (2017).

4. Herber, J. et al. Click Chemistry-mediated Biotinylation Reveals a Function for the Protease BACE1 in Modulating the Neuronal Surface Glycoproteome. Mol. Cell. Proteomics 17, 1487–1501 (2018).

5. Wollscheid, B. et al. Mass-spectrometric identification and relative quantification of N-linked cell surface glycoproteins. Nat. Biotechnol. 27, 378–386 (2009).

6. Bausch-Fluck, D. et al. A mass spectrometric-derived cell surface protein atlas. PLoS One 10, e0121314 (2015).

7. Boheler, K. R. et al. A human pluripotent stem cell surface N-glycoproteome resource reveals markers, extracellular epitopes, and drug targets. Stem cell reports 3, 185–203 (2014).

8. Post, H. et al. Robust, Sensitive, and Automated Phosphopeptide Enrichment Optimized for Low Sample Amounts Applied to Primary Hippocampal Neurons. J. Proteome Res. (2016). doi:10.1021/acs.jproteome.6b00753

9. Chen, J., Shah, P. & Zhang, H. Solid phase extraction of N-linked glycopeptides using hydrazide tip. Anal. Chem. 85, 10670–10674 (2013).

10. Furlan, C. et al. Miniaturised interaction proteomics on a microfluidic platform with ultra-low input requirements. Nat. Commun. 10, 1525 (2019).

11. Crisalli, P. & Kool, E. T. Water-soluble organocatalysts for hydrazone and oxime formation. J. Org. Chem. 78, 1184–1189 (2013).

12. Sobotzki, N. et al. HATRIC-based identification of receptors for orphan ligands. Nat. Commun. 9, 1519 (2018).

13. Gillet, L. C. et al. Targeted data extraction of the MS/MS spectra generated by data-independent acquisition: a new concept for consistent and accurate proteome analysis. Mol. Cell. Proteomics 11, O111.016717 (2012).

14. Bruderer, R. et al. Optimization of Experimental Parameters in Data-Independent Mass Spectrometry Significantly Increases Depth and Reproducibility of Results. Mol. Cell. Proteomics (2017). doi:10.1074/mcp.RA117.000314

15. Bausch-Fluck, D. et al. The in silico human surfaceome. Proceedings of the National Academy of Sciences 201808790 (2018).

16. Heng, T. S. P., Painter, M. W. & Immunological Genome Project Consortium. The Immunological Genome Project: networks of gene expression in immune cells. Nat. Immunol. 9, 1091–1094 (2008).

17. Liu, Y., Beyer, A. & Aebersold, R. On the Dependency of Cellular Protein Levels on mRNA Abundance. Cell 165, 535–550 (2016).

18. Lundberg, E. & Borner, G. H. H. Spatial proteomics: a powerful discovery tool for cell biology. Nat. Rev. Mol. Cell Biol. (2019). doi:10.1038/s41580-018-0094-y

19. Udeshi, N. D. et al. Antibodies to biotin enable large-scale detection of biotinylation sites on proteins. Nat. Methods (2017). doi:10.1038/nmeth.4465

20. Bruderer, R. et al. Extending the limits of quantitative proteome profiling with data-independent acquisition and application to acetaminophen-treated three-dimensional liver microtissues. Mol. Cell. Proteomics 14, 1400–1410 (2015).

21. Choi, M. et al. MSstats: an R package for statistical analysis of quantitative mass spectrometry-based proteomic experiments. Bioinformatics 30, 2524–2526 (2014).

